# Critical transitions in malaria transmission models are consistently generated by superinfection

**DOI:** 10.1101/355420

**Authors:** David Alonso, Andy Dobson, Mercedes Pascual

## Abstract

The history of infectious disease modelling essentially begins with the papers by Ross on malaria [1–5]. These models assume that the dynamics of malaria can most simply be characterized by two equations that describe the prevalence of malaria in the human and mosquito hosts. This structure has formed the central core of models for malaria and most other vector-borne diseases for the last century with occasional additions acknowledging important aetiological details. We partially add to this tradition by describing a malaria model that provides for vital dynamics in the vector and the possibility of super-infection in the human host; reinfection of asymptomatic hosts before they have cleared a prior infection. These key features of malaria aetiology create the potential for break points in the prevalence of infected hosts, sudden transitions that seem to characterize malaria’s response to control in different locations. We show that this potential for critical transitions is a general and underappreciated feature of any model for vector borne diseases with incomplete immunity and asymptomatic patients, including the canonical Ross-McDonald model. Ignoring these details of the host’s immune response to infection can potentially lead to serious misunderstanding in the interpretation of malaria distribution patterns and the design of control schemes for other vector-borne diseases.

## Introduction

Critical transitions occur when natural systems drastically shift from one state to another, they are currently receiving considerable attention in ecology, geophysics, hydrology and economics [6]. In epidemiology, critical transitions are of relevance to the emergence of new pathogens and the re-emergence of old ones. They may also be central to the within-host dynamics particularly when pathogens evolve or mutate in ways that allow them to escape control by the immune system. Critical transitions often underlie and potentially enhance (or undermine) attempts to control and eliminate infectious pathogens. Following an intervention, the trajectory of the host pathogen systems may cross a critical transition where pathogen prevalence drops to apparent eradication, the robustness of which is strongly determined by the structure of the transition. Tipping points associated with the coexistence of alternative equilibria are of particular interest, as small changes in a driving parameter can lead to large shifts from low to high levels of prevalence (or *versa vice*). Continuous external pressure on critical transmission parameters, or seasonal variation in vector abundance, can also lead to hysteresis, whereby a delayed response of the system would effectively keep it trapped longer in either the endemic or disease-free state.

Evidence for the existence of alternative steady-states in infectious disease dynamics remains limited [7–9], although they have been proposed as a possible explanation for the observation that malaria often fails to re-invade local regions that have achieved elimination. One potentially important pre-condition for the existence of alternative steady states in malaria is superinfection; the infection of a single host by concurrent multiple strains of the pathogen. Malaria infections are not fully immunizing, and multiplicity of infection is common in endemic regions as a consequence of additional infectious bites by the vector before the host has cleared a prior infection. In endemic regions, a large fraction of the human population carries the malaria parasite asymptomatically in non-apparent infections that can contribute to transmission. Under these conditions, significant levels of superinfection can create a positive feedback within the intensity of transmission that has the potential to generate multiple alternative equilibria and associated tipping points.

A large number of model formulations have been proposed to capture the complexity of malaria epidemiology [10, 11]. The consequences of repeated infectious bites have been represented in models of re-infection by a formulation that only allows a host to re-acquire infection after it has cleared a previous exposure. Two such recent models suggest that this might cause malaria dynamics to exhibit alternative steady-states [12, 13]. Here, we develop a complementary, but more general, analysis that provides a formulation for superinfection that explicitly allows infections to occur concurrently without interfering with each other. We initially present a semi-analytical approach to identify alternative equilibria in models for vector-transmitted diseases. We then apply these methods to a vector-borne disease model that was used to understand the origins of environmentally driven fluctuations of malaria in epidemic regions [14]. We then broadly consider superinfection in a series of hierarchical formulations of decreasing complexity (commencing with the original Ross-MacDonald equations). We demonstrate that irrespective of the details, **superinfection consistently creates tipping points** that can generate hysteresis in responses to control efforts (as well as seasonal variation in vector abundance). We argue that because models for re-infection and superinfection effectively bracket a continuum of different assumptions about within-host dynamics with multiple concomitant infections, then models that include these vital details of malaria biology should consistently exhibit tipping points in their response to control regardless of model formulation. Models that fail to include these effects may be misleading, or of limited utility when used to examine transitions towards low rates of transmission in response to control of vectors or of the pathogen.

## The model

The model is formulated as a system of ordinary differential equations that describe the transmission of the disease between the mosquito vector and the human host, it differs from the standard Ross-McDonald model by explicitly considering the vital dynamics of the two populations. Figure 1 presents the flow diagram of the model whose equations are described in detail in the Supplementary Material. The mosquito population is sub-divided into larval and adult stages, and adult individuals can be susceptible, infected or infectious. The human population has a constant size and deaths are assumed to exactly balance the birth rate; infected hosts are subdivided into two classes to differentiate individuals whose clinical infection leads to treatment from those whose natural recovery leads to the acquisition of immunity. These two classes map to symptomatic and asymptomatic infections respectively, the latter are crucially important as clinical infections typically represent only a small fraction of the total incidence in endemic regions [15–17]. Molineax, and Thomas [18], and Aron and May [19] scale the effective recovery rates, *r* by transmission intensity (the rate of infectious bites per human). Under the assumptions that infectious bites arrive at a constant rate, that the individual infections within a host proceed independently, and last a constant period, 1/*r*_0_, Dietz, Molineax, and Thomas [18] derived the following expression for the effective per capita recovery rate,

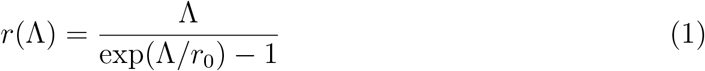

where Λ denotes the rate of infectious bites per human, and *r*_0_, is the basal recovery rate when disease transmission is very low (more precisely, in the limit of the infectious mosquito population tending to zero). Thus, the higher the rate of infectious bites per human host, Λ, the slower the disease clearance rate. When this rate Λ is measured per year, it is usually called the entomological inoculation rate (EIR). Finally, we assume that once individuals recover naturally from malaria (including the superinfection state with multiple concurrent infections), they enter a refractory period during which they are transiently immune and cannot re-acquire infection until they return to the susceptible class. Those who received treatment as the result of symptomatic infection move back to the susceptible class.

**Figure 1:**
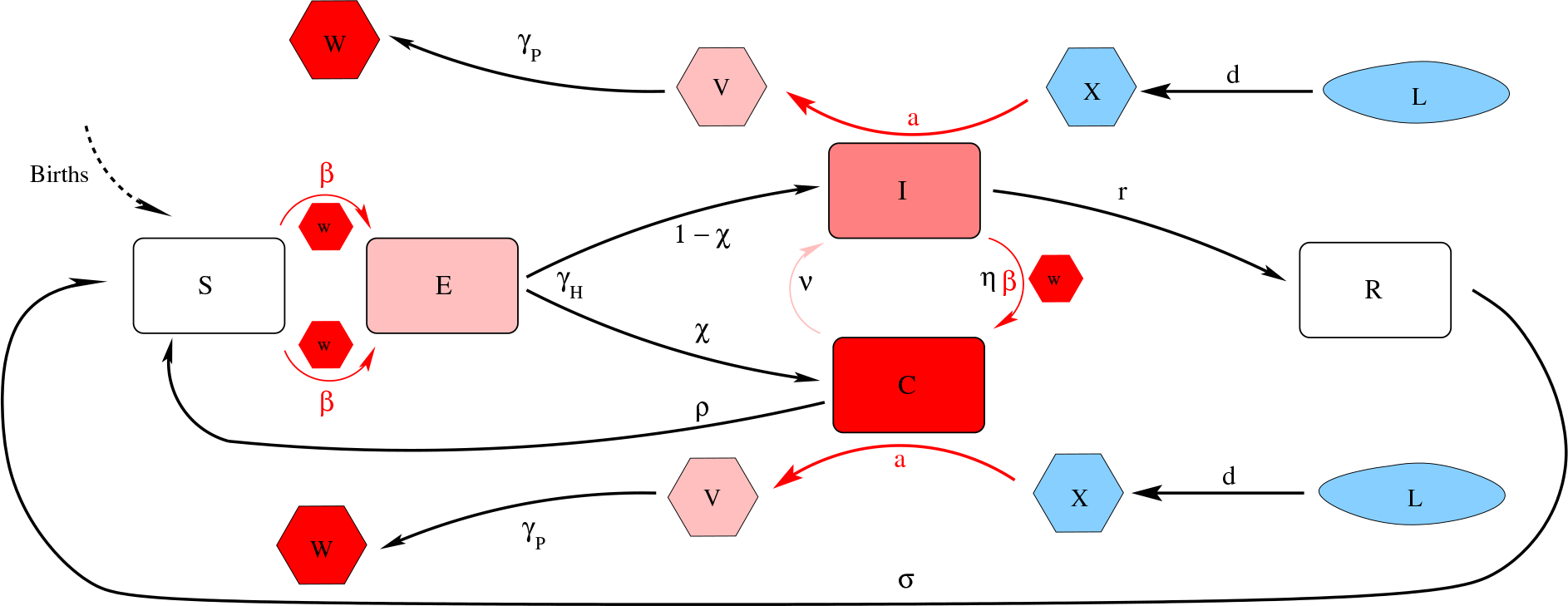
The human-mosquito SECIR-LXVW coupled model. The human stages of infection ‘SECIR’ are depicted in the central part of the figure progressing from S to E to I from left to right. The different stages of the mosquito population, from larva (*L*) to infectious adult mosquito (*W*), are drawn in the upper and bottom of the figure progressing from uninfected L, larvae, and adults (blue) to incubating (pink) and infected adults (red), from right to left (LXVW). Per capita death rates (respectively *δ* and *δ*_*M*_, and *δ*_*L*_ in the equations) affect all mosquito and human classes, and for simplicity are not included in the diagram.

The couplings between human and mosquito dynamics are given by the force of infection *β*, the per capita rate at which a susceptible human contracts the disease from infectious bites, and by the per capita rate *λ* at which adult mosquitoes acquire the pathogen from infectious humans. Under reasonable assumptions [20], these two rates can be written as

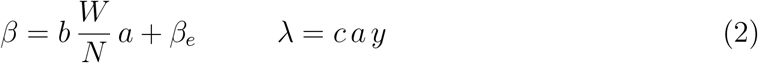

where *b* and *c* are the respective probabilities that humans develop infection once bitten and that mosquitoes acquire the parasite from biting an infectious human host. Moreover, 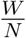 *a* is the number of infectious bites per human host (see definition of Λ above), *β*_*e*_ is an external force of infection, and *y* is the fraction of infectious humans (see Supplementary Material for details).

Our model can be seen as an extension of the classical formulation introduced by Ross and MacDonald [21] (RM) (see also [10, 20, 22] for a review). Although the original RM model has given rise to a multiplicity of malaria models [11] spanning different degrees of complexity that range from delayed ODEs to SIR (Susceptible-Infected-Recovered)-like structures, we show that these models are all related to the RM formulation through a set of sequentially simplifying assumptions. In particular, all converge to the RM model which is based on two strong simplifications: (1) constant mosquito (*M*) and human populations (*N*), and (2) only two classes (susceptible and infectious) representing vectors and humans (see Supplementary Material, for details). We start with the analysis of alternative steady states in the full model of Fig 1, and end with the consideration of their more general existence in a suite of models of decreasing complexity. Model parameter space has been explored between maximum and minimum values (around two different parameter combinations — A and B, see Table S1 in Supplementary Material) that provided a good fit to data in a previous study from an epidemic region [14].

## Results

To identify the stationary points of the coupled system we present a semi-analytical method that consists of first finding the equilibria of the two submodels, namely the expression for infectious mosquitoes as a function of infectious humans and vice versa. The fixed points are then obtained by calculating the intersection of these two curves (see Fig 2). The generality and feasibility of this method rest on the linearity of the human and mosquito submodels when considered separately. This means that, for a given number of infectious mosquitoes (*W*\*), the human submodel becomes a linear ODE system. And, likewise, for a given fraction of infectious humans (*y*), the mosquito submodel becomes a linear ODE system as well. We summarize the main steps of the approach below (full details are given in the Supplementary Material). Once the stationary points have been fully characterized, we can map several quantities of interest onto the parameter space, including the model reproductive number *R*_0_, the expected number of secondary cases produced by an initial single infection in a completely susceptible population, and the dominant period, *T*_0_, associated with the damped oscillations in the transients towards stationarity.

**Figure 2:**
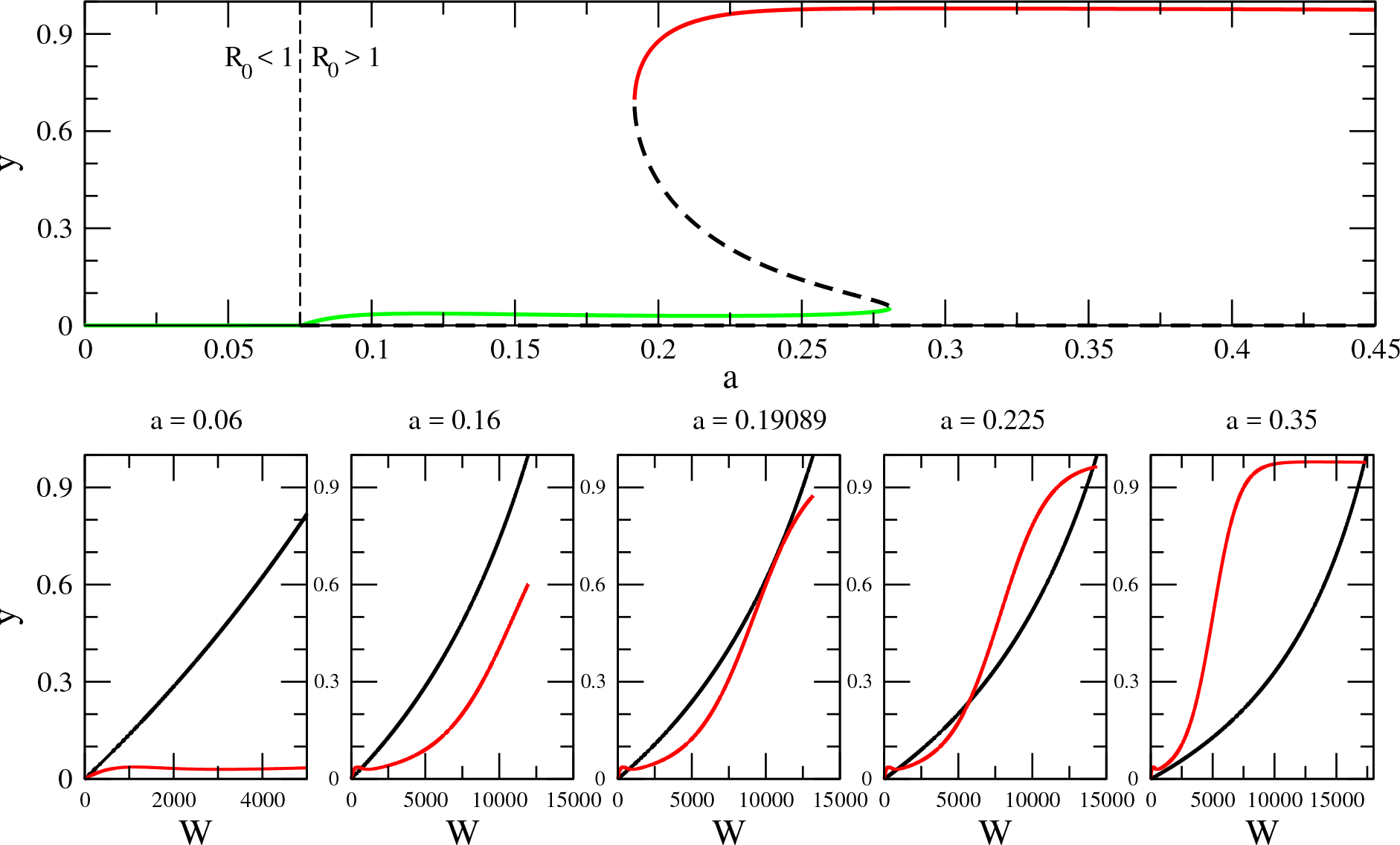
Relationship between saddle-node bifurcation and mosquito biting rates. When biting rates are low, the curves intersect at zero and malaria fails to establish resulting in a disease-free equilibrium (*R*_0_ < 1). As the biting rate *a* increases, the two curves intersect initially at one single point (a=0.16) corresponding to a stable endemic equilibrium. When the biting rate crosses a critical value (*a*_*C*_ = 0.19089), a tangential bifurcation arises. Further increases of the biting rate end up producing two alternative stable states corresponding to the coexistence of two endemic equilibria characterized high and low malaria incidence (*a*_*C*_ < *a* < 0.28). Eventually, only the upper equilibrium is left (*a* >0.28). The full bifurcation diagram shown in the upper plot is calculated with parameter combination A (but with *β*_*e*_ = 0, see Table S1). It represents a cross section of Fig. 4b following the horizontal broken line shown there (but here starting at *a* = 0 rather than at *a* = 0.15) for incremental increases in biting rate

### Mosquito submodel: stationary points

To obtain the expression for infectious mosquitoes at equilibrium, one first needs to calculate the fixed point for total mosquito abundance. It is easy to show that the mosquito population model has only one globally stable point (Supplementary Information):

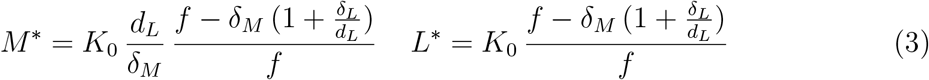

A single condition controls when this point is a feasible stable point for the dynamics:

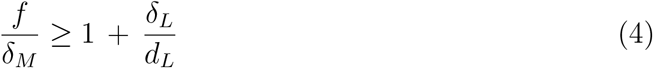

Biologically, this condition means that the mosquito population will be locally maintained in a given area when the number of larvae recruited per female during their adult average life time compensates for the larval losses during their development stage. This threshold condition arises in the full expression for *R*_0_ (see Eq S49 and S51) because only if *M*\* is positive, *R*_0_ is non-negative, so it is weil-defined.

### Human-mosquito model: stationary points

The equilibrium number of infectious mosquitoes *W*\* is derived by solving the mosquito ODE subsystem at equilibrium when the fraction of infected humans *y* is considered as a fixed parameter. We obtain the curve

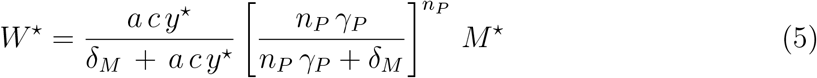

A 2nd curve is then calculated by solving the human submodel at equilibrium for a given level of infectious mosquito population, this makes *β* a constant parameter. This curve for the stationary fraction of infectious humans is given by

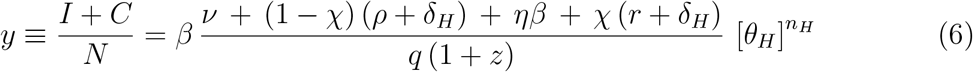

where the total force of infection, *β*, depends on the number of infectious mosquitoes per human, with *q* and *z* standing for composite parameters. Full expressions for these parameters and fixed points are given in the Supplementary Material.

### Coexistence of alternative stable states

Fig. 2 shows that the intersection of the curves represented by Eqs (5) and (6) can produce more that one fixed point. When intersections are present, the corresponding stable points can be computationally calculated with a cobweb procedure (see Fig S2). Although this procedure cannot converge to the intermediate unstable point, this can be calculated with a standard bisection method [23]. The pair (*W*\*, *y*\*) defines the stationary state and the full solution can then be unfolded from it (see Supplementary Material for details). Fig 2 also shows that as the biting rate *a* increases, the system undergoes several bifurcations. Each of them represents the emergence of a new stationary point. The first one corresponds to a transcritical bifurcation [24] and represents the transition from a free-disease situation (*R*_0_ < 1) to an endemic stable equilibrium (*R*_0_ > 0). This is the typical bifurcation of infectious disease models found in SIS, SIR, and SEIR systems [25]. The second one corresponds to a saddle node bifurcation (also called a tangential or fold bifurcation). The tangential intersection of the two curves defines a critical biting rate (*a*_*C*_ = 0.19089) and for *a* > *a*_*C*_, the sudden appearance of a pair of resting points, a saddle node and a second stable point with higher disease incidence. The first stable point corresponding to a lower incidence endemic equilibrium remains. As a result, two basins of attraction coexist, each consisting of initial conditions that lead to one of the two alternative stable states, separated by the existence, of an intermediate unstable state. Thus, for the same parameter values and depending on initial conditions, the system can end up in one of two possible stable equilibria for high and low disease incidence.

The emergence of the first transition (from disease-free to endemic equilibrium) can be characterized by calculating the reproductive number *R*_0_. The transcritical bifurcation corresponds to *R*_0_ = 1 (see vertical broken line in the upper panel of Fig 2). We calculated this quantity as the dominant eigenvalue of the next generation matrix [26] (see Supplementary Material). In addition, we mapped *R*_0_ onto the sub-parameter space given by the plane defined by the carrying capacity of the vector *K*_0_ and its biting rate *a* (see Fig 3a). These two parameters determine the intensity of transmission. The expression underlying *R*_0_ is the same for the model with and without superinfection since this quantity depends only on the basal rates *r*_0_ and *σ*_0_ (see Supplementary Material, Eqs. S49 and S51). In addition, for the regions where damped oscillations occur, the dominant period of those oscillations can be also calculated. In the region of the parameter space we studied (see Table S1, supplementary material), we found dominant periods between about 2.5 to 11 years (see Fig 3b).

**Figure 3:**
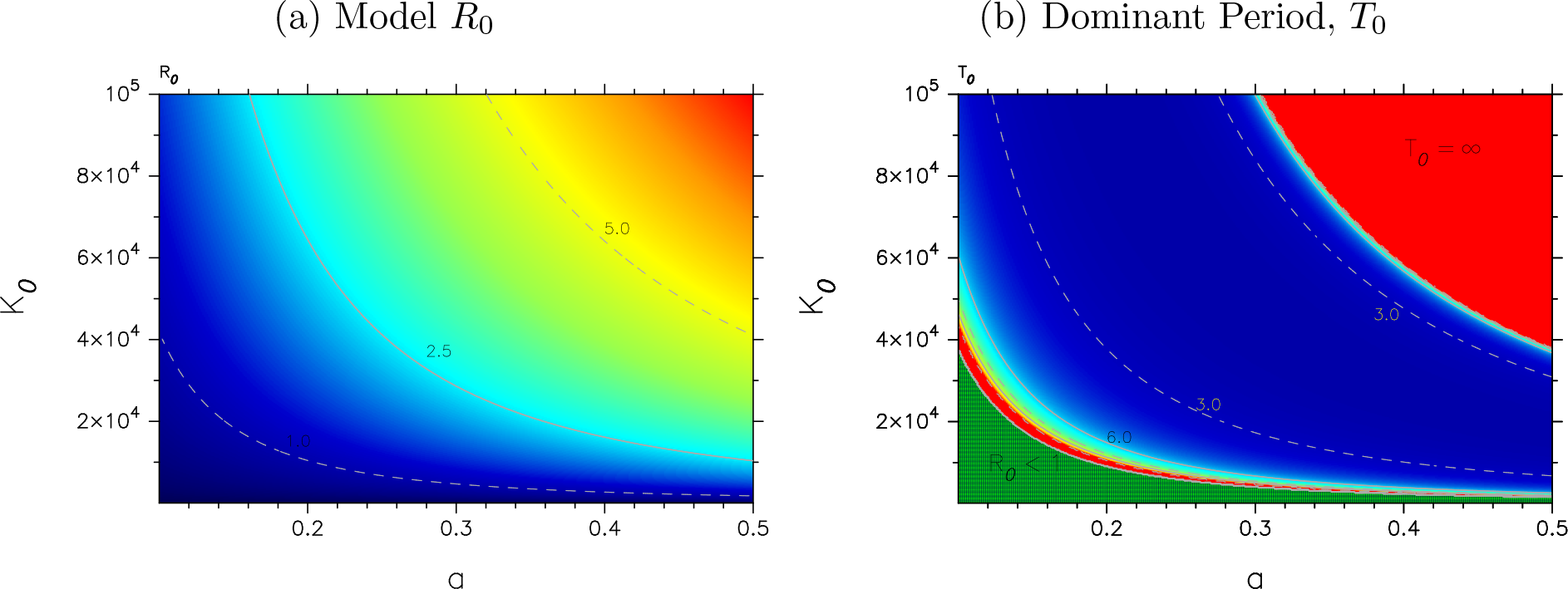
(a) The relationship between *R*_0_, mosquito carrying capacity *K*_0_, and biting rate *a*. (All other parameter values are those of parameter set B (with *β*_*e*_ = 0, Table S1, Supplementary Material). (b) The corresponding relationship between the dominant period of the damped epidemic oscillations, *T*_0_ in the same plane of parameter values (see Fig S1, Supplementary Material). In the region where bi-stability is found, the natural period corresponds to damped oscillation to the lower stable equilibrium. (This period is computed from the complex eigenvalues of the Jacobian evaluated at this lower equilibrium). Red and green areas have the same interpretation as in Fig 4. Selected contour lines representing periods of 3 and 6 years are also shown for guidance.

Identifying the different regimes of the system requires us to determine the local stability of the different fixed points. The Jacobian matrix *J*\* is evaluated at the corresponding fixed points and its eigenvalues are obtained as the roots *λ* of the characteristic polynomial:

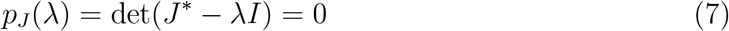

where *I* is the identity matrix. Because this polynomial can have as many roots as the dimension of the dynamical system, a fixed point is asymptotically locally stable, if and only if its associated eigenvalues *λ*_*i*_ have all strictly negative real parts. In that case, if the imaginary part of the dominant eigenvalue is not null, damped oscillations are observed in the approach to the fixed point. We determined different properties of the stationary state, such as the possibility of coexisting fixed points, parameter-induced bifurcations, and the presence of endogenous (damped) oscillations. The resulting possible dynamical regimes are mapped onto different subregions of the parameter space in Fig. 4. Comparison of the upper and lower panels shows that the coexistence of alternative steady states critically depends on including superinfection in the model.

**Figure 4:**
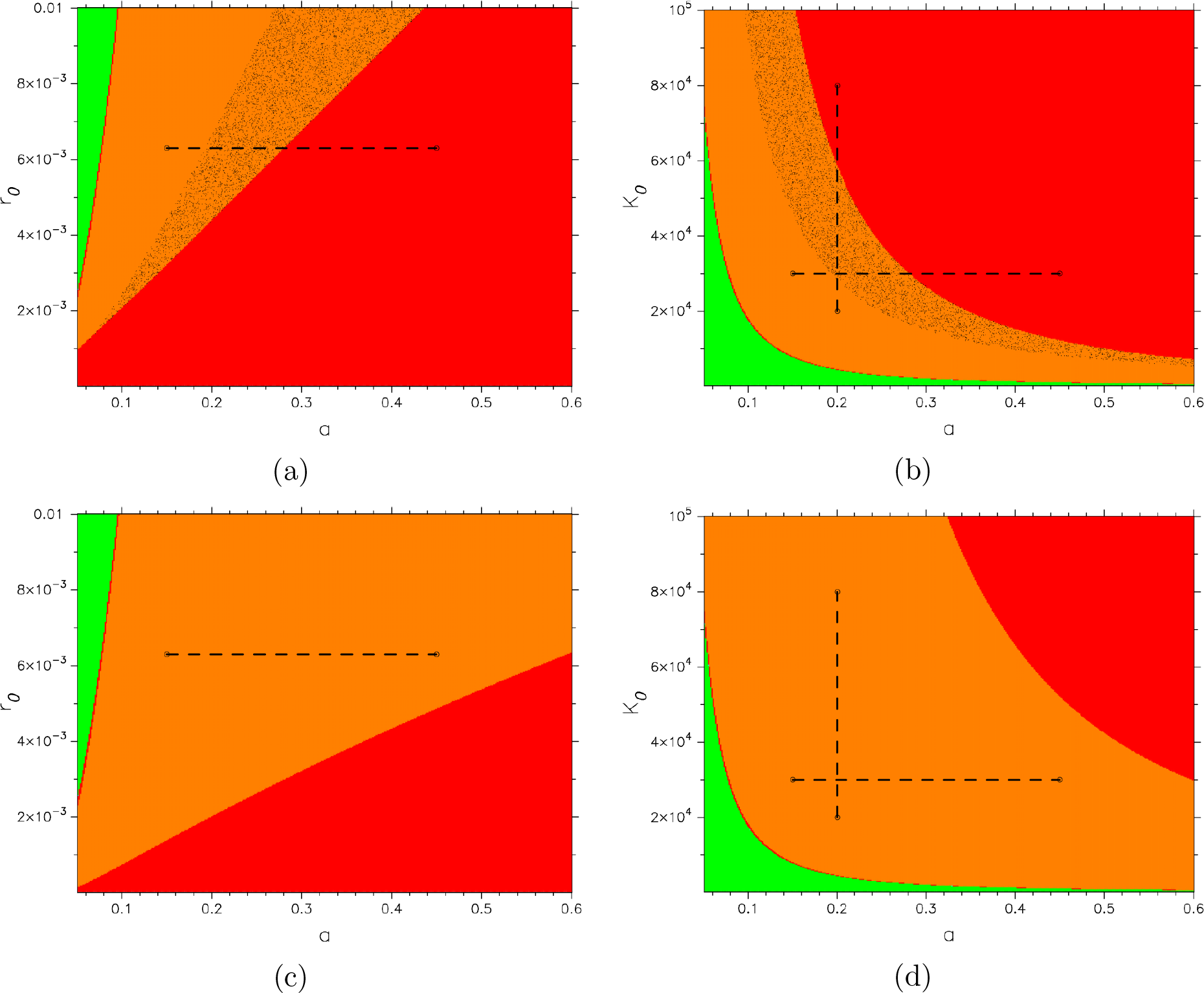
Dynamic Regimes. Relationship between key parameters in the model that govern the emergence of bifurcations. In all cases the x-axis depicts the biting rate *a* and the y-axis either the basal recovery rate *r*_0_ (left) or the carrying capacity *K*_0_ (right). The colored regions delineate different domains of dynamics behavior: non-trivial disease equilibria that are locally stable and approached with endogenous damped oscillations are obtained for parameters in the orange region; within this region, two stable equlibria coexist in the shaded area (panels A and B). Non-oscillating stable endemic equilibria occur only in the red region. The region of parameter space where the system exhibits global disease-free equilibria is shown in green in panel B. The fraction of infectious humans exhibit a bifurcation in the section illustrated by the broken lines in the upper panels. The lines cross a region where a single stable equilibrium gives rise to two coexisting stable equilibria as the value of each of the bifurcation parameters increases (see also panel C of Fig. 5 and panel D of Fig S). The lower panels illustrate phase diagram for the same model structure and parameter combinations but without superinfection (Solution A, with *β*_*e*_ = 0, from Table S1, Supplementary Material). This corresponds to the more widely used model that assumes constant rates of recovery (*r* = *r*_0_) and loss of immunity (*σ* = *σ*_0_); clearly these simplifications fail to capture the tipping points and multiple equilibria seen the upper figures (see Supplementary Material).

Our procedure to calculate fixed points is completely general and can be applied to any coupled vector-human model. It is independent of specific assumptions about the force of infection, including the existence of an external source of infection, as well as other possible ways in which both the mosquito and human submodels might be defined, as long as both of these are linear ODE systems when considered separately. In the Supplementary Material, we add further realism and apply the approach to a system where the exponential distribution of disease incubation times in both human and mosquito submodels has been replaced by the more general (gamma) distribution, which effectively reduces to a fixed duration of infection or incubation [27]. The stationary solution reduces to the case illustrated in Fig 4 when *n_H_* = *n_V_* = 1 (see also Table S1).

### Hysteresis

Coexistence of stable equilibria can underlie sharp transitions between levels of disease prevalence if a sufficient perturbation is applied to the system to move it from one basin of attraction to the other. The responses to more gradual changes in parameters can instead lead however to hysteresis [6], this is of key interest to malaria control, since the dynamic memory of the system can delay responses and allow the persistence of endemicity beyond the bifurcation point at which we would expect elimination to occur. A related further possible effect is an asymmetry in the temporal trajectories from endemicity to elimination, and from elimination to re-emergence. These hysteresis effects are illustrated in Fig 5 for slow changes in the biting rate *a*; these might occur during the transition between dry and rainy seasons. We consider first changes from an initial biting rate *a* = 0.1 at *t*_0_, to a value of 0.5 at the intermediate time *t*_*i*_, and then back to its initial value at time *t*_1_. The response of the system is illustrated for the fraction of infectious humans (*y* = (*C* + *I*)/*N*), a measure of disease incidence, where *N* is total human population (panels B and C). The response of the system to a linear increase of the biting rate is a non-linear increase in disease incidence (see Fig 5), this can be represented as a hysteresis cycle on the corresponding bifurcation diagrams (Panel C). In fact, extensive initial increases in *a* leave disease levels almost unchanged up to certain threshold. In addition, there is clear asymmetry in the two opposite control trajectories: the identical linear trajectory back to the initial low levels of the biting rate does not return the system to the same initial levels of disease incidence (see Fig 5). Beyond the threshold, the system has been pushed towards a higher incidence state from which it is hard to return. Although decreasing *a* back to its initial low values would eventually lead the system to settle down at the initial low incidence equilibrium determined by imported infections (for *β*_*e*_ ≠ 0), the transient trajectory to this final state can be very long. Although everything else remains constant, the amplitude of hysteresis cycles depends on how fast the driving parameter changes. The same behaviors can be illustrated by measuring system response in terms of the entomological infectious rate (Fig. S3, Supplementary Material).

**Figure 5:**
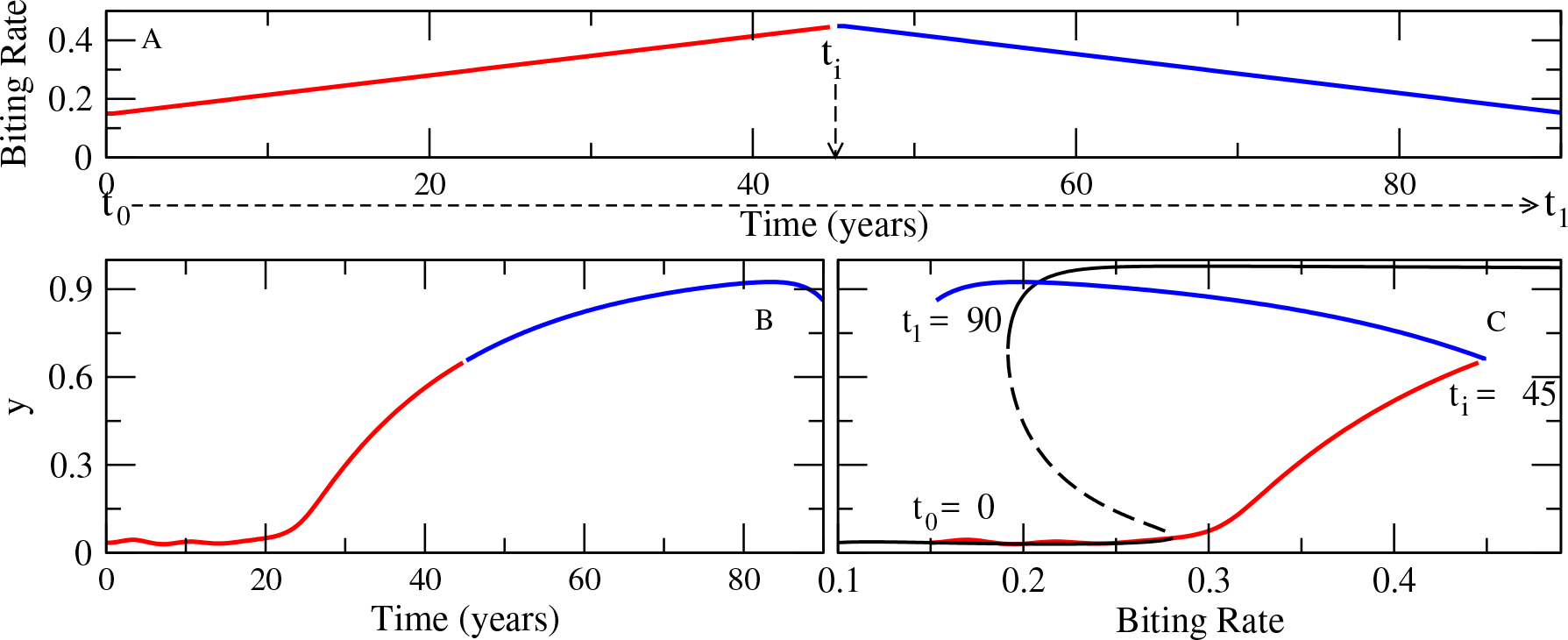
Hysteresis Cycles. Response of the fraction of infectious humans to a slowly increasing biting rate *a*, from low to high values and back (Panel A). Other parameter values are given by set A of Table S1. We plot the time evolution of the system when considering superinfection. The blue and red curves correspond to dynamics with a respective biting rate increasing and decreasing as a linear function of time. The diagrams correspond to changes in *a* respresented by the horizontal broken line of the upper left panel in Fig 4. In the presence of the slowing-down effect, as biting rates go from low to high and back to low values in a linear way (panel A), disease incidence (*y*) shows a highly non-linear response (see panel B), which can also be represented as a hysteresis cycle in the bifurcation diagrams (see panel C).

### Model Robustness

In order to quantify how the presence of bi-stability in malaria models depends on their degree of complexity, we evaluated the relative size of the region of parameter space where this behavior arises for different formulations (Fig 6). We specified 16 different human-mosquito coupled ODE models of increasing realism. The models are labeled according to the number of classes considered for the human and mosquito populations: four possibilities for the former (*SE*_*n*_*CIR*, *SE*_*n*_*IR*, *SE*_*n*_*I* and *SI*), and four for the latter (*LXV*_*n*_*W*, *X_K_V*_*n*_*W*, *XV*_*n*_*W*, and *XW*). Without exhausting possible variations, this scheme results in a total of 16 different combinations, which are labeled as *SE*_*n*_*CIR* − *LXV*_*n*_*W*, *SE*_*n*_*CIR*−*X*_*K*_*V*_*n*_*W* (see Supplementary Material for a detailed description of each model). When a model increases in complexity by the addition of new model parameters (encoding particular processes), the fraction of parameter space leading to a given dynamical regime can increase, remain unchanged, or decrease. The fraction of parameter space where a given regime is found is calculated by randomly drawing parameter sets within minimum and maximum values (as given in Table S1), and evaluating the fraction of those draws leading to a given dynamic regime. Fig 6 compares models from the perspective of the relative size of regions with endemic equilibria or bi-stability. The exercise suggests that the coexistence of alternative stable states is robust to model simplification. This is shown here by representing the probability of bistability as model complexity decreases, from the *SECIR* − *LXV*_*n*_*W*, a model with 21 parameters, to the *SI* − *LXV*_*n*_*W*, a model with only 13 parameters (see Fig 6b). Even the simplest mosquito-human coupled model (*SI* − *XW*), with only 8 parameters, corresponding to the RM formulation, exhibits this feature as long as the positive feedback introduced by the slowing down of the human recovery rate under repeated infectious bites is maintained (see Fig S4 and Supplementary Material). For each of these model combinations, the removal of the slowing down effect in the recovery rate that is produced by superinfection eliminates bi-stability (as shown in Fig 4, panels C and D). We also show the effect of one added parametric dimension, the introduction of an external force of infection, in models that combine the mosquito subcomponent *LXV*_*n*_*W* with human submodels of decreasing model complexity (Fig 6b). These’open models (*β*_*e*_ ≠ 0)’ show a relative decrease in the fraction of the parameter space showing bi-stability, mainly due to the linearizing effect of external infections.

**Figure 6:**
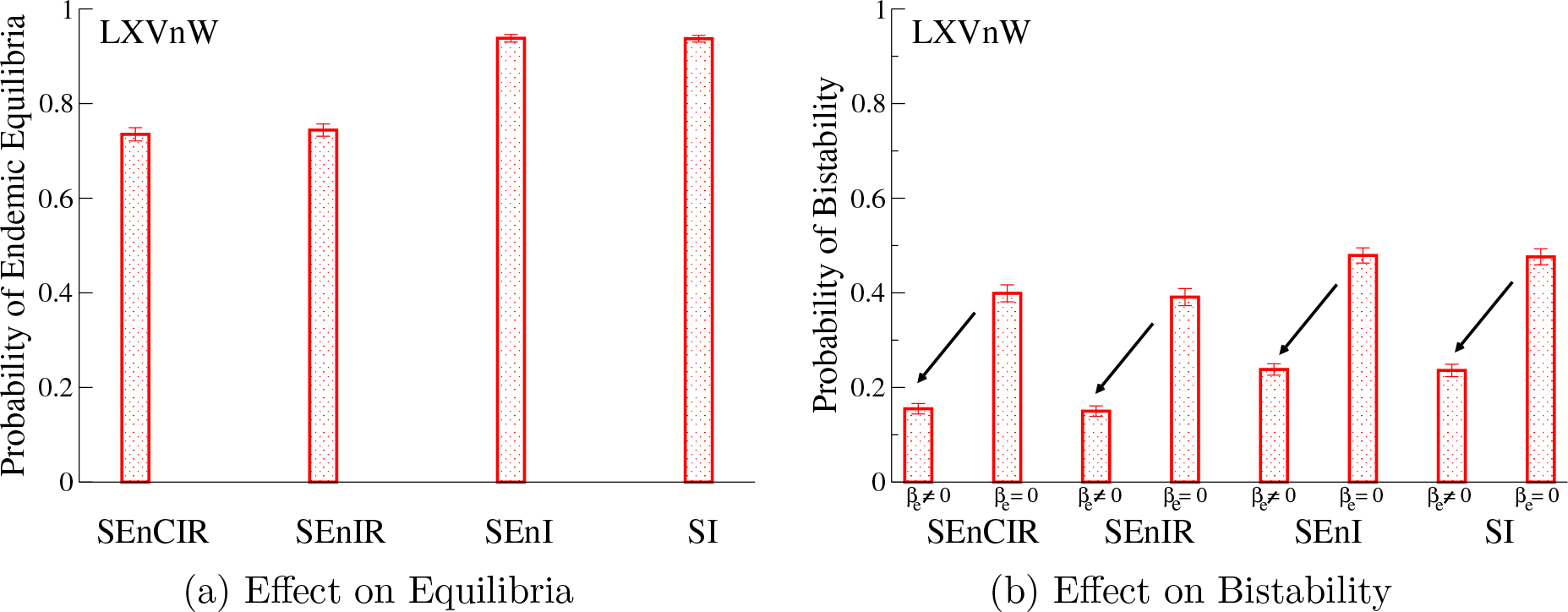
Effect of model complexity resulting from the addition of human and mosquito classes (and the corresponding increase in the number of parameters), on the fraction of the full parameter space corresponding to either the existence of a single endemic equilibrium (a) or the presence of bi-stability (b).

In sum, superinfection, this is, the slowing-down effect on recovery rates as infectious bites increase, is a necessary condition for the presence of bi-stability. However, we showed that bistability only arises in certain areas of the parameter space. Under superinfection, further refined necessary conditions for a given parameter can be derived provided others parameter values remain constant. The presence of bi-stability can be studied by comparing the slopes of the curves represented in Fig 2 at the origin for *W* = 0. By using this strategy, for instance, for the model *SI* − *XW*, we derived a necessary condition for the onset of bi-stability in terms of *b*, the probability of infection of a susceptible human upon receiving an infectious bite. This condition *b*_*c*_ < *b* < 0.5 indicates that *b* has to be lower than 50% but higher than a certain critical threshold *b*_*c*_ determined by the other model parameters (see derivation and threshold value in Supplementary Material).

## Discussion

Immunity to malaria clearly differs from that in standard epidemiological models assuming full protection upon recovery (classical SIR models). It is widely accepted that naturally acquired immunity involves diverse immunity responses to the different phases of the parasite within humans; whereas pre-erythrocytic and sexual stages of the life cycle of the parasite are poorly immunogenetic, merozoites are mainly targeted by the immune system [28]. Multiplicity of infection (MOI) and the exhorbitant antigenic diversity of the *Plasmodium* in endemic regions complicates the acquisition of immunity in ways that can only be captured phenomenologically in standard transmission models. More realistic representations than the one presented here include the explicit consideration of MOI and strain variation in individual-based models (e.g. [29, 30]), and the age of hosts in age-structured (partial differential equations) models [31]. Stage-structured models could also be used to represent increasing levels of immunity acquired through repeated exposure, beyond the two-stage models considered so far [12, 31].

Our work demonstrates that inclusion of superinfection in malaria models, not only determines the lengthening of infectious periods [18], but is a key factor responsible for the coexistence of multiple stationary states, and the possibility of nonlinear regime shifts. Our model specifically considers the slowing down effect of transmission intensity (rate of arrival of infectious bites) on the recovery rate (*r*) and therefore, the duration of infection. A second effect is on the lengthening of the duration of immunity (through *σ*). Only the former is needed for the occurrence of alternative steady states. A model including solely the effect on immunity duration lacks this behavior, whereas one including the effect of superinfection on infection duration exhibits the same behaviors and transitions illustrated in our results. This observation finds a clear explanation in the contrasting effects of these two assumptions: a lengthening of infectious periods produces a positive feedback on infection duration and transmission rate. With longer infections, a higher number of vectors acquire the parasite, increasing the rate of infectious bites, and concomitantly slowing further the recovery rate. As illustrated for a wide range of ecological systems, positive feedback is the key ingredient for the strong nonlinearities that underlie regime shifts and associated alternative equilibria [6]. In contrast, a deceleration of the loss of immunity slows down the return of resistant hosts to the susceptible class, and the resulting higher number of immune individuals decreases the number of infected vectors. This negative feedback on transmission intensity is incapable of producing by itself coexistence of alternative stable states.

Our derivation for the implications of superinfection for a critical transition is robust to consideration of a wide range of malaria models spanning different levels of complexity. Even the simple RM model exhibits bi-stability when the recovery rate slows down with increasing transmission intensity.

Previous malaria models showing bi-stability differ from ours in that they assume re-infection, and therefore only allow a host to re-acquire the parasite after complete clearance of infection and its return to the susceptible state [12, 13]. When our findings are taken together with those of these earlier models, they underscore the broad generality of critical transitions in malaria models regardless of specific biological details on multiple infections and within-host dynamics. This generality follows from the observation that re-infection and superinfection effectively bracket a continuum of possible assumptions on the outcome of repeated exposure to infectious bites. In the former, repeated infections completely interfere with each other; in the latter, they do not ‘see’ each other at all. For endemic regions, multiplicity of infections is the rule and not the exception. Although it cannot be captured by only either re-infection or superinfection assumptions. clearly, reality must fall somewhere in between depending on the particular representation of within-host dynamics. Neither complete interference (re-infection) nor a complete lack of interference (superinfection) are likely.

A different kind of mechanism independent from repeated exposure has been proposed for alternative steady-states in malaria dynamics. This mechanism combines density-dependent biting rates and disease-induced mortality [7]. This mechanism assumes that increasing disease levels would decrease total human population through disease-induced mortality, which then makes biting rates higher since they depend on the density of infectious mosquitoes per human, which, in turn, would rise infection levels in the human population. However, we believe this positive feedback is less important in nature; it should also work in conjunction with the main mechanism of repeated exposure, either through re-infection or superinfection, described here.

The possibility of nonlinear thresholds in the mosquito-malaria system has several important implications. Small changes in parameters (for instance, biting rate *a* or mosquitoes’ carrying capacity *K*) can give rise to large changes in incidence. Control efforts may see no progressive decrease of incidence until a sudden effect finally occurs. In the opposite direction, elimination states may be more robust than in the case of standard transcritical transitions. Also, the progressive relaxation of control efforts in endemic regions could generate sudden transitions from low to high incidence. Finally, concerning variability, sudden shifts from low to large fluctuations in incidence may follow in epidemic regions from environmental condition such as temperature warming driving the system across a critical threshold [14].

An important next step is the confrontation of these kinds of dynamics to data from surveillance efforts across changing control and environmental conditions (e.g. [31]). On-going developments on early-warning systems related to critical transitions [32–34], and the prediction of the distance and kind of bifurcation based on the monitoring of a few large perturbations, should be brought together with sustained surveillance efforts over time and space and theoretical findings on the behavior of malaria models.

